# Dynamics of sensorimotor plasticity during spatial finger augmentation

**DOI:** 10.1101/2025.07.01.662639

**Authors:** Dominika Radziun, Siebe Geurts, Valeria C. Peviani, Luke E. Miller

## Abstract

How does the brain integrate artificial body extensions into its somatosensory representations? Prior work shows that the body and technology are represented simultaneously, but little is known about how these representations evolve dynamically across different sensorimotor interaction phases. To fill this gap, we investigated the dynamics of somatosensory plasticity using a custom-built wearable finger-extension device that elongated users’ fingers by 10 cm, spatially augmenting their reachable workspace. Across four time points, before, during (pre-and post-use), and after device wear, participants completed a high-density proprioceptive mapping task to measure representations of the biological and artificial fingers. We observed three distinct phases of plasticity. First, wearing the finger-extension device led to a contraction of the perceived length of the biological finger. Second, after active use, the represented lengths of the biological and artificial fingers stretched significantly, an effect absent when participants trained with a non-augmenting control device. Third, a post-removal aftereffect on the biological finger representation was observed. These results demonstrate that wearable finger extensions that provide spatial augmentation are rapidly integrated into body representations, with dynamic proprioceptive adjustments shaped by structural and functional properties of the device. This work advances our understanding of how the sensorimotor system accommodates artificial extensions and highlights the potential for body-augmenting technologies to be intuitively integrated within the sensorimotor system.

## Introduction

The human body is remarkably adaptable, capable of integrating artificial enhancements that extend its physical capabilities (Miller and Farnè, 2024). This includes traditional tools (e.g., rakes) that operate outside the body as well as body augmentation devices that directly interface with the user, including supernumerary robotic limbs (SRLs), extra digits, and articulated hand or arm exoskeletons (Prattichizzo et al., 2021; Yang et al., 2021). Decades of research has found that tools influence the user’s body representations and their implementation in the brain (Maravita and Iriki, 2004; Martel et al., 2016). In contrast, relatively little is known about how body augmentation devices themselves are represented by the user’s somatosensory system. Furthermore, most studies have used pre-post experimental designs, offering a limited temporal window into the effects of technology. In our study, we aimed to address these gaps by investigating how representations of the body and technology dynamically evolve across different interaction phases.

Both tools and augmentation devices drive changes in the user’s body representations (Berti and Frassinetti, 2000; Eden et al., 2022; Gallivan et al., 2013; Holmes and Spence, 2004; Iriki et al., 1996; Longo and Serino, 2012; Miller et al., 2019b; Peviani et al., 2024). Tools alter their user’s spatial tactile perception and motor kinematics in ways consistent with stretching the sensorimotor representation of the user’s arm (Bruno et al., 2019; Canzoneri et al., 2013; Cardinali et al., 2011, 2009; Galigani et al., 2020; Garbarini et al., 2015; Martel et al., 2019; Miller et al., 2017, 2014; Najafabadi et al., 2023; Romano et al., 2019; Sposito et al., 2012). Beyond tools, a broad class of articulated augmentations—SRLs mounted at the shoulder or waist, exo-digits, and multi-DOF hand exoskeletons—expand the user’s motor control abilities (Abdi et al., 2016; Hussain et al., 2016; Maekawa et al., 2020; Mooney and Herr, 2016; Penaloza and Nishio, 2018; Prattichizzo et al., 2021; Shafti et al., 2021; Umezawa et al., 2022; Yang et al., 2021). Controlling an extra robotic finger can modify body representation in sensorimotor cortices (Kieliba et al., 2021), aligning with evidence that tool use modulates somatosensory processing and induces plasticity in body representations, particularly in secondary somatosensory and posterior parietal cortices (Miller et al., 2019b).

The human nervous system can use proprioceptive feedback to extract the spatial extent of hand-held tools (Debats et al., 2017; Fabio et al., 2024; Miller et al., 2018; Peviani et al., 2024; Solomon and Turvey, 1988; Themelis et al., 2025). This suggests that the sensorimotor system constructs spatial representations of tools using similar computations as for the body (Miller et al., 2023). Evidence that the somatosensory system also represents the spatial extent of body augmentation technology is limited (Dowdall et al., 2025; Nishida et al., 2020). Furthermore, it is an open question whether—like the body—representations of technology undergo spatial plasticity (e.g., stretching) during motor learning—similar dynamics would be expected if body and technology were represented similarly.

The majority of the above studies assessed plasticity only before and after use, typically once the device was removed from the body. How body representations evolve while the technology is being worn and used has received little attention. Furthermore, the extent to which representations of biological and artificial segments are concurrently represented and interact remains unclear. Therefore, identifying the influence of technological augmentation at all distinct stages—from the initial wearing of the technology to its use and subsequent removal—is crucial for understanding the dynamics of plasticity.

To characterize the temporal dynamics of plasticity, we developed a passive, non-articulated wearable device that rigidly attaches to the fingertips and increases the effective length of each finger by 10 cm, thereby spatially augmenting their reachable workspace. Participants completed a proprioceptive localization task (Peviani et al., 2024) over four stages: before and after wearing the device, and before and after performing motor training tasks that required functional use of the device. This high-density proprioceptive localization task measured how accurately participants could localize different parts of their finger and the artificial finger in space, allowing us to quantify changes in body representations at different time points. By sampling these distinct interaction phases and jointly mapping biological and artificial finger segments, our approach significantly extends prior demonstrations of body-representation plasticity and sensorimotor remapping with a time-resolved account of how a spatial augmentation of the hand is integrated.

We observed three time-specific heterogenous effects of the finger-extension device: (i) an initial compression of the user’s biological finger representation upon wearing the device, (ii) a stretch of proprioceptive representations of both the user’s biological and artificial fingers that was driven by training to use the augmentative device—an effect that was absent after using a non-augmenting device, and (iii) a stretch of the user’s finger representation that persisted after the device was removed and was likely driven by persistent exposure to wearing the finger extensions. Together, these findings offer theoretically important insight into how the brain dynamically integrates artificial extensions into the body representations.

## Results

### Wearable finger extensions

To enable proprioceptive mapping of both the biological and artificial finger segments, we constructed a custom passive wearable finger-extension device that rigidly coupled to the user’s distal phalanges and elongated each finger by 10 cm (Fig. 1A). The finger-extension device was 3D-printed and designed to form a collinear extension of the biological finger with no additional joints or actuation. To enable efficient object manipulation, it was comprised of a rigid internal spine mimicking the bone and a soft silicone outer layer allowing for compliant contact with objects (Fig. S1). Unlike a typical tool (e.g., pliers), the movements of the biological finger joints were directly transmitted to the translation of the finger extensions. Therefore, the biological and artificial finger segments formed a single kinematic chain that could be sensed by the somatosensory system. This design feature was critical for the goals of our experiment, as it allowed us to map a participant’s proprioceptive geometry continuously along both segments within a shared reference frame. A structurally matched non-augmenting version (1.4 cm extension) served as the control device in Experiment 1B (Fig. 1A).

**Figure 1.**
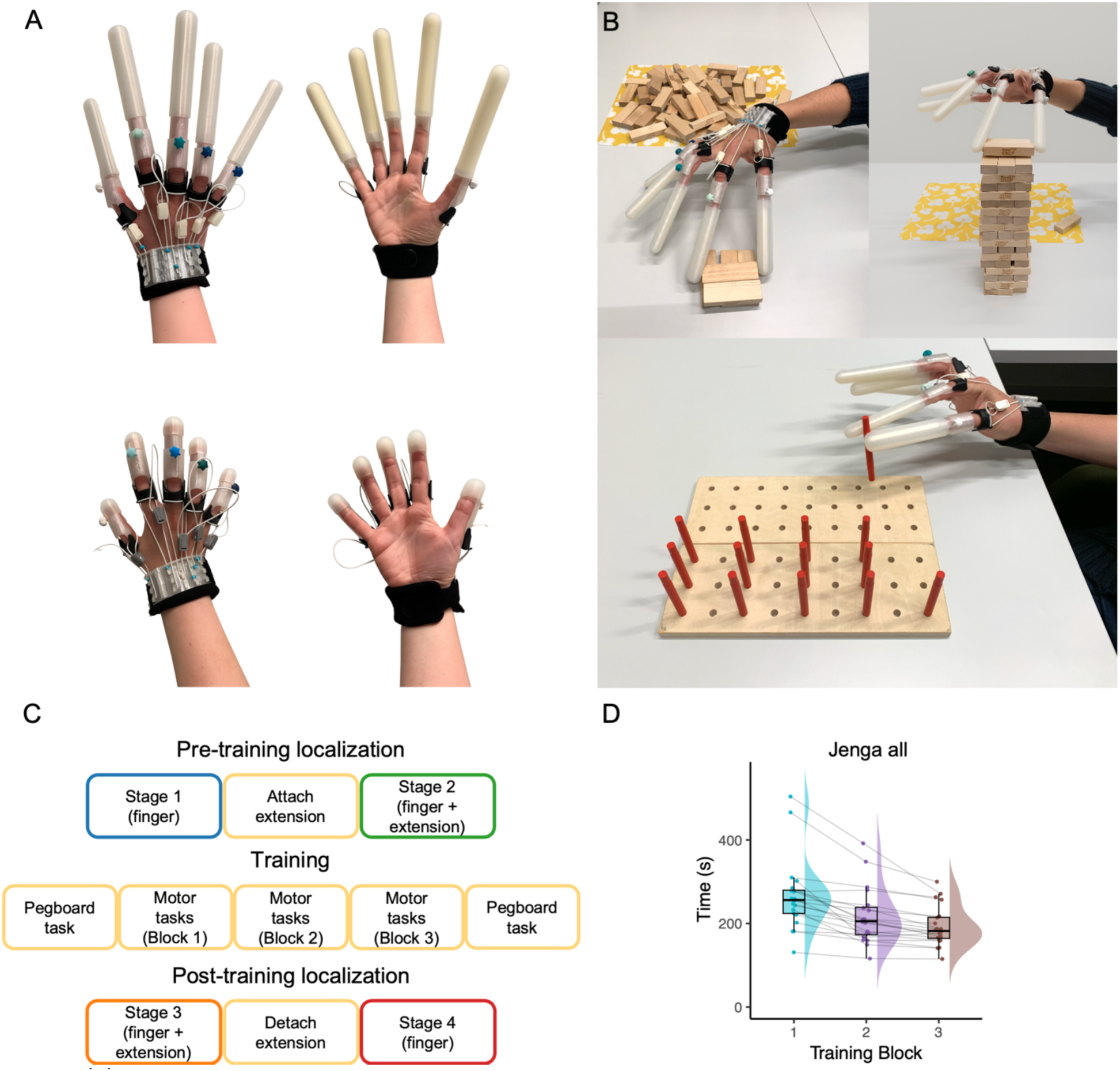
(A) The experimental, wearable finger-extension device with 10 cm finger extensions and the control device without the extensions. (B) Example of the Jenga stacking training task. Participants wore the finger-extension device and were asked to stack a full Jenga tower; task performance was measured by the time taken to complete the tower. (C) Outline of the experimental design. (D) Group data showing improvement in performance (reduction in completion time) across training blocks.

A variety of motor tasks were employed to train participants to use these devices (Fig. 1B; Supplementary Video 1). These included sorting colored blocks, stacking a Jenga tower, and sorting pegs into holes; the latter task required fine-grained manipulation and was used as our outcome measure of motor learning.

### Experiment 1A: Introduction to task protocols and device familiarization

The goal of our study was to elucidate the temporal dynamics of somatosensory representational plasticity during body augmentation. To this end, we explored the effects of finger length augmentation on somatosensory finger representation. Participants (n=18) performed a high-density proprioceptive localization task in which they used their left hand to indicate on a touchscreen where they felt specific target locations on their right index finger, which remained hidden under the screen. The task involved repeated localization of seven equally spaced points along the length of the finger (and later, also the artificial finger), enabling fine-grained assessment of perceived spatial structure.

This task was performed at various time points relative to learning to use the finger-extension device (Fig. 1C): before wearing the device (stage 1); while wearing the device, before using it (stage 2); while wearing the device, after using it (stage 3); and after the device was removed (stage 4). In all stages, we mapped the participant’s representation of their biological index finger. In stages 2 and 3, we also concurrently mapped the participant’s artificial finger representation. Reported *p*-values are Bonferroni-corrected to account for five planned comparisons.

### Wearing the finger extensions contracts perceived biological finger length

Participants initially localized seven proprioceptive targets—equally spaced along the midline of the right index finger from the base to the fingertip—by touching the corresponding locations on a touchscreen (stage 1). Each target was judged 20 times and the perceived location for each target was the mean of those repetitions (see also Fig. S3A for the target layout). We then used linear regression to characterize their biological finger representation, modelling their localization judgments as a function of actual target location. This analysis revealed that participants’ representation closely tracked actual finger geometry, with regression slopes moderately above one (mean±sem: 1.16±0.10, 95% CI [0.95, 1.39]).

Participants were then fit with the finger extensions, providing them a short period of visual and proprioceptive feedback about their mechanics and geometry (around 1-2 minutes; see Supplementary Video 2). They then performed the proprioceptive localization task while wearing the device (stage 2). The mapping now included 14 targets: seven on the biological finger and seven on the artificial finger segment (Fig. S3B). Importantly, the task mapped both the biological and artificial fingers simultaneously (bio+extension).

When wearing the finger extensions, the combined bio-extension map was geometrically accurate, spanning the true length of the combined surface (mean true length: 17.40 cm; mean estimated length: 18.90 cm; paired *t*-test of map versus true length: *t*(17) = 1.24, *p =* .23, BF_10_ = 0.47). How this space was partitioned was approximately proportional across surfaces. Specifically, the scaling of the biological and artificial finger representations did not differ significantly (bio slope mean±sem: 0.98±0.07; extension slope mean±sem: 1.04±0.07; two-tailed paired t-test: *t*(17) = −0.8, *p* = 1.00, BF_10_ = 0.34), with the biological finger accounting for 41.2% of the combined representation (actual: 42.5%). Comparing the biological finger slopes with the map measured during stage 1 suggested an apparent contraction in represented length after attaching the finger extensions (16.27% reduction; *t*(17) = 2.34, *p* = .030 [uncorrected], *p* = 0.16 after correction, BF_10_ = 2.09; Fig. 2A). This effect was influenced by a single outlier participant; when this individual was excluded, the contraction became stronger and statistically robust (20.48% reduction; *t*(16) = 3.67, *p* = 0.002 [uncorrected], *p* = 0.01 after correction, BF_10_ = 20.24). Given that a comparable contraction was also observed in Experiment 1b (see below), we interpret this as evidence for a reliable reduction in represented biological finger length following attachment of the finger extensions.

**Figure 2.**
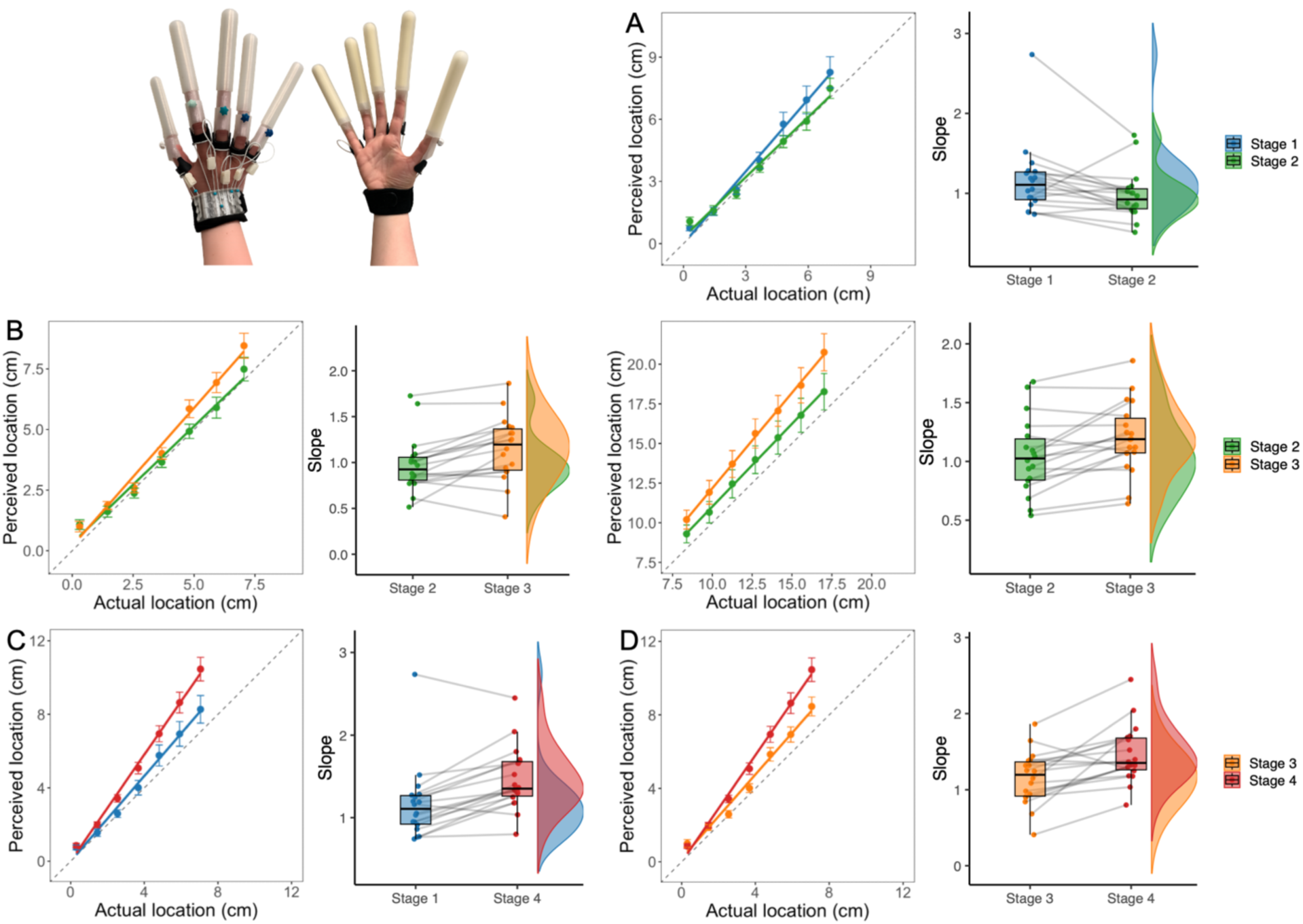
Proprioceptive judgments across stages of augmentation with wearable finger-extension device. (A) Comparison of proprioceptive maps from stage 1 (baseline) and stage 2 (initial device fitting). (B) Left: Comparison of biological finger segment between stage 2 and stage 3 (after training). Right: Comparison of artificial finger segment between the same stages. (C) Comparison of biological finger representation from stage 1 to stage 4 (post-device removal). (D) Comparison of biological finger representation from stage 3 (with device) to stage 4 (after removal). In all line plots, perceived location (y-axis) is plotted against actual target location (x-axis). Error bars represent standard error of the mean. The dashed line indicates the identity line (slope = 1). Raincloud plots show individual participant slopes in each stage. The following figure is formatted in the same fashion.

### Active use of the finger extensions leads to proprioceptive “stretching” of biological and artificial finger segments

After stage 2, participants completed a 30-45 minute training period to familiarize themselves with the finger-extension device (Fig. 1B). Training performance improved significantly over time, indicating a clear learning effect and increased proficiency with the device (Fig. 1D; Fig. S4). Detailed analyses of individual tasks consistently showed significant improvements across sessions, confirming this effect. Full statistical results and task-specific analyses are provided in the Supplementary Information.

Following training, participants again completed the bio+extension finger localization task (stage 3). Based on previous research with tool use (Bruno et al., 2019; Canzoneri et al., 2013; Cardinali et al., 2011, 2009; Galigani et al., 2020; Garbarini et al., 2015; Martel et al., 2019; Miller et al., 2017, 2014; Najafabadi et al., 2023; Romano et al., 2019; Sposito et al., 2012), we hypothesized that training would stretch the user’s finger representation. We found that participants’ internal mapping of the combined finger (bio+extension) was no longer geometrically accurate (true length: 17.40 cm; estimated length: 21.16 cm; paired t-test: *t*(17) = 3.08, *p =* .007, BF_10_ = 7.27). This served as an initial confirmation of our hypothesis that training with the finger extensions led to a representational stretch.

We then examined how proprioception of both the biological finger and the artificial finger segment changed following functional augmentation. The perceived length of the biological finger increased following training (slope mean±sem: 1.15±0.08, 95% CI [0.98, 1.32]; 17.93% increase; *t*(17) = 3.67, *p* = .009 BF_10_ = 23.45; Fig. 2B), demonstrating that finger augmentation stretched the user’s finger representation. Interestingly, we identified a similar effect of representational stretch for user’s representation of the artificial finger (slope mean±sem: 1.20±0.07, 95% CI [1.06, 1.35]; length change: 15.35% increase; *t*(17) = 3.77, *p* = .008, BF_10_ = 26.06; Fig. 2B). Though stretched, the relative portion of the representation devoted to the finger remained stable (finger proportion 41.16%; paired t-test on proportions before and after training: *t*(17) = 0.02, *p* = 1, BF_10_ = 0.24). Together, these findings indicate that proprioceptive representations of both the biological and artificial segments were stretched following training with the finger extensions while preserving their relative allocation within the combined map.

### Proprioceptive aftereffects emerge following device exposure

After completing the bio+extension localization task (stage 3), participants performed the localization task again with the finger-extension device removed (stage 4). This allowed us to examine whether representational changes to the biological finger remained at this immediate, post-removal measurement. We did this by comparing the final representation length, measured in stage 4 (slope mean±sem: 1.46±0.09, 95% CI [1.28, 1.65]), with two previous stages: stage 1 and stage 3. Regardless of which set of stages we compared the final mapping to, we observed a similar a modulation in the length of the participant’s biological finger representation: an 25.35% increase compared to stage 1 (Fig. 2C; paired t-test: *t*(17) = −5.15, *p* < .001, BF_10_ = 344.52); and an increase of 26.94% compared to stage 3 (Fig. 2D; *t*(17) = –3.85, *p* = .006, BF_10_ = 30.18).

### Experiment 1B: A non-augmenting device does not stretch the user’s proprioceptive representations

We next sought to determine whether the observed training-driven representational stretch (i.e., stage 2 versus 3) was specific to training with a spatial finger-augmentation device. To this end, the same eighteen participants took part in the above experimental procedure with one change: the training was done using a control device that did not meaningfully elongate the fingers (1.4 cm extension; Fig. 1A). Note that all proprioceptive mapping still included the finger-extension device as in Experiment 1A. This allowed us to hold constant the mapped surface (the long extension) and time spent on the task; therefore, any stage 2 to stage 3 change could be attributed specifically to the presence of spatial augmentation during training rather than to proprioceptive mapping alone.

As in Experiment 1A, the combined bio-extension map obtained during stage 2 (initial localization with the finger-extension device) was overall geometrically accurate (true length: 17.40 cm; estimated length: 19.35 cm; paired t-test of map versus true length: *t*(17) = −1.41, *p* = .18, BF_10_ = 0.57). The biological and artificial finger segments were represented with similar scaling (bio slope mean±sem: 1.04±0.09; extension slope mean±sem: 1.07±0.08; paired t-test: *t*(17) = −0.44, *p* = 1, BF_10_ = 0.27), indicating no significant difference between the two segments. However, comparing stage 1 (pre-device) to stage 2, we observed that attaching the finger extensions significantly contracted the represented length of the biological finger (16.15% reduction; *t*(17) = 4.02, *p* = .004, BF_10_ = 41.57; Fig. 3A). The magnitude of this compression was similar to Exp. 1A (*t*(17) = 0.18, p = 0.91, BF_10_ = 0.245; Fig. 4A). This suggests that, despite similar scaling factors of the two segments, wearing the finger extensions alone led to a measurable compression of the biological finger representation.

**Figure 3.**
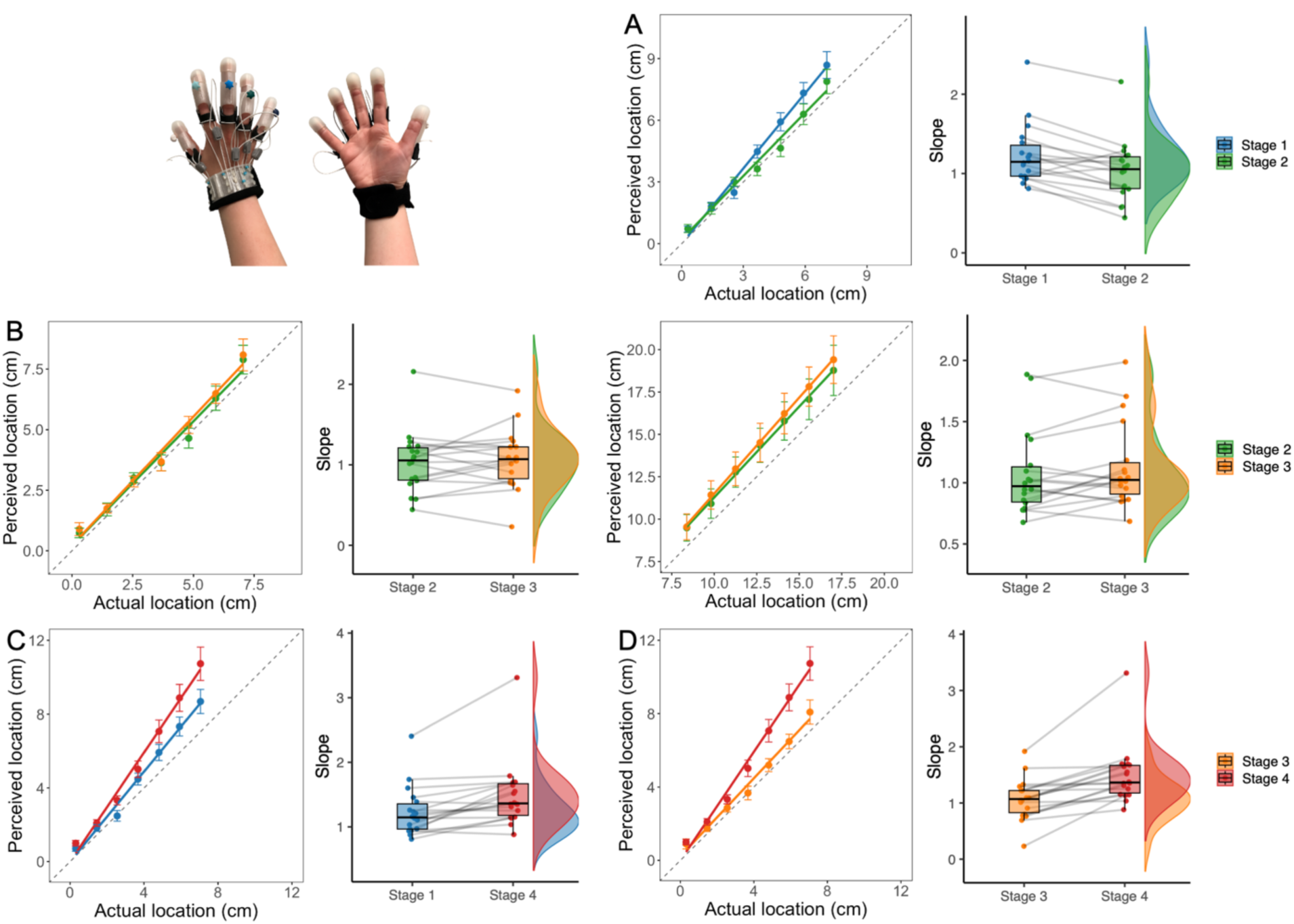
Proprioceptive judgments across stages with training performed using the control device. (A) Comparison of proprioceptive maps from stage 1 (baseline) and stage 2 (initial finger-extension device fitting). (B) Left: Comparison of biological finger segment between stage 2 and stage 3 (after training with control device). Right: Comparison of artificial finger segment between the same stages. (C) Comparison of biological finger representation from stage 1 to stage 4 (post-device removal). (D) Comparison of biological finger representation from stage 3 (after training with control device) to stage 4 (after removal). In all line plots, perceived location (y-axis) is plotted against actual target location (x-axis). Error bars represent standard error of the mean. The dashed line indicates the identity line (slope = 1). Raincloud plots show individual participant slopes in each stage.

**Figure 4.**
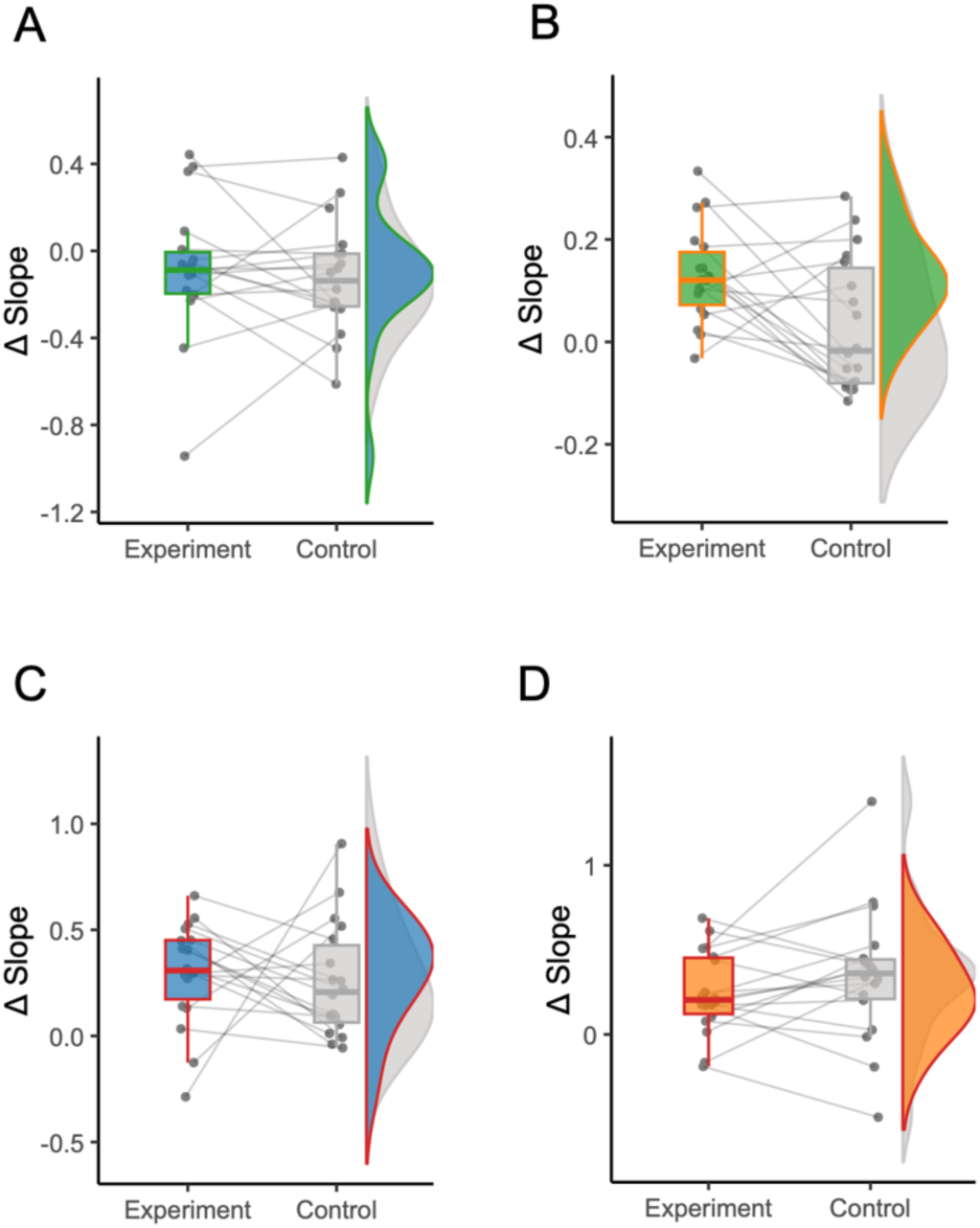
Direct comparison of representational changes across training devices. Difference scores in proprioceptive localization slopes across stages, shown separately for the spatially augmenting finger-extension condition (Experiment 1A) and the non-augmenting control condition (Experiment 1B). (A) Difference between Stage 1 and Stage 2. (B) Difference between Stage 2 and Stage 3. (C) Difference between Stage 1 and Stage 4. (D) Difference between Stage 3 and Stage 4. In all panels, each boxplot shows the group distribution for one device condition, with individual participant values overlaid.

We next determined the effect of using the control device on the biological and artificial finger representations. Participants were immediately fit with the finger-extension device after their training with the control device. Unlike in Experiment 1A, we did not observe changes in the represented length of the biological (mean±sem: 1.07±0.08, 95% CI [0.89, 1.25]; *t*(17) = −0.70, *p* = 1.00, BF_10_ = 0.30; Fig. 3B) or artificial finger (mean±sem: 1.13±0.08, 95% CI [0.96, 1.3]; *t*(17) = −1.68, *p* = .56, BF_10_ = 0.78; Fig. 3B). This pattern suggests that stretch of somatosensory representations is specific to the active use of the spatially augmenting finger extensions, and not simply due to the attachment or observation of them. Furthermore, although participants improved on several training tasks in this control session (e.g., pegboard, sorting, Jenga timing; see Supplementary Information), these practice gains did not translate into proprioceptive stretching at stage 3 when the augmenting device was fitted, indicating that the training-dependent expansion observed in Experiment 1A is augmentation-specific.

To directly test whether training with the spatially augmenting finger-extension device produced greater representational stretching than training with the non-augmenting control, we ran a 2 x 2 repeated-measures ANOVA on the combined bio+extension slope with within-subject factors Stage (2 vs. 3) and Device (finger-extension device [Exp. 1A] vs. control device [Exp. 1B]). There was no main effect of Device (*F*(1,17) = 0.18, *p* = .674), a main effect of Stage (*F*(1,17) = 17.03, *p* < .001), and, critically, a Stage x Device interaction (*F*(1,17) = 7.05, *p* = .017; Fig. 4B). The interaction indicates that training with the finger extensions selectively increased the combined representation length from Stage 2 to 3 relative to training with the non-augmenting control.

Interestingly, we replicated the effect of removing the finger extensions in stage 4. We observed a similar a modulation in the length of the participant’s biological finger representation: a 20.9% increase compared to stage 1 (Fig. 3C; paired t-test: *t*(17)= −4.05, *p* = .004, BF_10_ = 44.16); an increase of 39.29% compared to stage 3 (Fig. 3D; paired t-test: *t*(17)= −4.98, *p* = .001, BF_10_ = 248.43). Intriguingly, we did not observe a significant difference between the degree of change in Experiment 1A (*t*(17) = −0.36, *p* = .72, BF_10_ = 0.26; Fig. 4C-D). These results suggest that the proprioceptive aftereffects measured in both experiments was not due to the training stage, but likely to other factors such as wearing the finger-extension device and using proprioceptive feedback of its geometry to perform the localization task. The distinct pattern of plasticity observed in both experiments after using the finger extensions (stage 2 versus 3) and removing them (stage 4) hints at distinct internal models for the hand and hand+extensions (see Discussion).

## Discussion

This study provides a detailed characterization of how the human somatosensory system dynamically adapts to a wearable device that spatially augments the reachable workspace of the fingers. Using a high-density proprioceptive localization task, we show that spatial representations of both the biological body and an artificial extension can be flexibly reshaped across time. Specifically, we identified three distinct phases of somatosensory plasticity: an initial contraction of biological finger representation upon wearing the device, a functional “stretching” of both the biological and artificial finger maps following training, and a post-removal aftereffect observed at our final measurement after the finger-extension device was removed. Importantly, the emergence of these changes following training demonstrates that humans can quickly form a somatosensory representation of the artificial fingers based on sensory feedback. While the initial contraction occurred with both the finger-extension and control devices, the functional stretching and its associated post-removal effect were specific to augmentation with an actively used finger-extension device that spatially augmented the user’s fingers. Together, our findings reveal how the somatosensory system integrates novel artificial extensions (Makin et al., 2017, 2023; Miller et al., 2019b).

We show that wearable finger extensions are an effective means for investigating the dynamics of plasticity in body representations. Unlike hand-held tools, these devices are directly interfaced with the body (Maravita and Iriki, 2004; Martel et al., 2016). Learning to use such a device introduces new demands on proprioceptive processing, requiring the integration of additional structure into the body’s representational framework (Raisamo et al., 2019). Here, we observed that training with a spatially augmenting finger-extension device led to measurable changes in how both biological and artificial finger segments were represented. These learning-driven changes were absent when participants used a control device that did not augment finger length, highlighting the specificity of functional engagement in driving plasticity (Biggio et al., 2020; Umiltà et al., 2008).

The initial act of wearing the finger extensions, even before use, triggered a reallocation of representational space, leading to a contraction in the represented length of the biological finger (i.e., an under-representation). This suggests that the somatosensory system responds immediately to the presence of an artificial augmentation, even in the absence of task-specific learning. While this early “miniaturization” recalls reports from studies with passive exoskeletons (e.g., HandMorph), in which participants appeared to judge objects as larger and perceived their grasping range as reduced on first wear (Nishida et al., 2020), those findings may partly reflect task-specific factors such as altered hand posture. Nonetheless, both results point to a rapid adjustment of body representation upon donning novel augmentative devices. These observations align with theories of neural resource allocation, in which new sensory inputs initially compete with existing representations for limited perceptual or cortical resources (Dominijanni et al., 2021). The contraction of the biological finger’s representation may reflect an adaptive mechanism, reallocating capacity toward processing novel sensorimotor structures (Makin et al., 2017; Miller et al., 2019b).

Following training, we observed that both the biological and artificial finger representations underwent a significant stretch, an effect selectively present after training with finger extensions and absent after control training. This parallel plasticity suggests that the brain may apply shared representational mechanisms to both body parts and technological extensions, reinforcing the idea that augmentation devices can be integrated into the sensorimotor system in a body-like manner (Makin et al., 2023, 2020; Miller et al., 2023, 2019a). The absence of such effect in the control condition emphasizes that this plasticity is not simply a response to attachment, but a specific outcome of using augmentation to achieve new functional outcomes.

One complementary way to viewing our results is via the concept of *internal models*, predictive representations of body dynamics that aid in perception and motor control (Scott, 2004; Shadmehr and Krakauer, 2008; Wolpert et al., 2001). Previous work suggests that internal models play a critical role in tool embodiment, allowing the brain to simulate and predict the outcomes of actions involving both the body and its extensions (Kilteni and Ehrsson, 2017; Miller et al., 2018). If a user forms biological+technological internal models, then motor learning should modify both similarly. Indeed, we found that the proportion of the overall representation dedicated to the biological and artificial finger segments remained constant before and after the observed plasticity. This suggests that the presence of a mechanism that globally stretches the representation of both surfaces, as predicted if the user learned a biological+extensions internal model. This internal model likely underlies the system’s ability to coordinate perception and action during interaction with the finger extensions (Imamizu et al., 2000; Miller et al., 2018).

This biological+extensions internal model appears to be newly learned during the experiment and is distinct from the user’s pre-existing internal model for their fingers. Notably, simply wearing the finger extensions led to a clear contraction of the biological finger representation, reflecting an immediate reallocation of representational space upon donning the device. This initial compression was followed by an opposite pattern once the device was removed. We observed a post-removal aftereffect—an increase in biological finger length perception following removal of the device—that occurred both when participants had actively used the finger extensions and when they had only trained with a control device. This suggests that modifying this finger representation did not depend on motor training or functional use of the extension, as was the case with the newly learned internal model. Instead, the pre-existing internal model could be modified with repeated sensory-motor engagement involved in the proprioceptive localization task itself. Importantly, this engagement occurred while participants were still wearing the device, but its perceptual consequences became apparent only once the finger extension was removed and the mapping returned to the biological finger alone. Together, the initial contraction and later expansion effects indicate that wearing the device alone induces immediate representational compression, whereas only active use of the augmenting device drives the subsequent stretching observed during training.

Despite providing detailed insights into somatosensory plasticity during spatial finger augmentation via a wearable device, our study has limitations that should be considered. The experiments were conducted under short-term, controlled conditions and did not include active sensory feedback from the device, limiting our understanding of how such plasticity unfolds in more ecologically valid or long-term contexts. Although proprioceptive localization provides a high-resolution behavioral readout of body representation, it offers limited leverage on how different sensory channels contribute to these effects. Systematically introducing tactile or force feedback could reveal how these inputs shape augmentation-related plasticity. Prior work suggests that even simple visuotactile pairing can support the embodiment of additional digits (Cadete and Longo, 2020), underscoring the potential of multisensory stimulation in shaping body representation. Moreover, recent reviews highlight that supplementary sensory feedback improves control performance and embodiment for supernumerary robotic limbs (Pinardi et al., 2023). Therefore, these results may provide a behavioral template for the design of next-generation exoskeletons and prosthetics that incorporate artificial sensing to simulate touch. Exploring how the brain integrates these richer feedback channels could help make the artificial extensions to be experienced more like biological body parts rather than external tools. To move toward practical applications, future studies should investigate how neural representations of artificial fingers evolve with extended use and more complex sensorimotor feedback. These directions are essential not only for understanding sensorimotor plasticity, but also for informing the non-invasive restoration of touch in assistive technologies (Nisky and Makin, 2024).

In sum, this work demonstrates that the human sensorimotor system is capable of rapidly and flexibly integrating wearable finger extensions that provide spatial augmentation into its spatial framework. Crucially, this integration is shaped by both the physical characteristics of the augmentation and the nature of the user’s interaction with it. By showing that proprioceptive representations can quickly adapt to artificial fingers, we provide empirical support for the development of intuitive, body-integrated assistive technologies. Understanding the neural and perceptual mechanisms by which the brain incorporates such devices has direct implications for neurorehabilitation, prosthetics, and the design of human-machine interfaces that feel like natural extensions of the self.

## Materials and Methods

### Participants

A total of 20 healthy right-handed participants were recruited (mean age = 27.2 years, range = 18–54; 13 women, 7 men), all of whom completed two experimental sessions on separate days. Two participants were excluded due to the use of undesired strategies to perform the task, resulting in a final sample of 18. One used their visible left hand to mirror the position of their unseen right hand, resulting in atypically high accuracy and low uncertainty. The other switched midway from proprioceptive localization to using a fixed visual reference point, producing a bimodal response distribution. All participants had normal or corrected-to-normal vision and no history of neurological or psychiatric conditions. Participants were asked not to have long nails on the day of the experiment, as this could interfere with the usage of the devices. The research was approved by the Ethics Committee of the Faculty of Social Sciences, Radboud University. Every participant gave informed consent to participate in the experiment. Participants were recruited through the SONA system of Radboud University and received a 30-euro voucher or course credits as compensation.

### Finger-extension devices

The devices used in this study were custom-built passive wearable devices designed to augment finger function while interfacing directly with the user’s hand (Fig. S1). The finger-extension device (Exp. 1A) elongated each finger by 10 cm while preserving the participant’s native finger-joint mobility. This device rigidly coupled to the distal phalanges using fingertip caps, such that the biological finger and its artificial finger segment formed a single collinear mechanical module. This design ensured that finger flexion and extension were transmitted directly to the artificial finger, without introducing additional joints or actuation. As a result, the device modified the reachable workspace of the hand without altering its joint kinematics. For the control condition, a structurally similar device with an effective elongation of only 1.4 cm was used (Exp. 1B).

#### Structure and materials

The structural frame of the finger-extension device was 3D-printed in PETG filament on Prusa MK3 and MK4 printers, with higher-wear joint components printed in PET. The outer surface of each artificial finger was cast in a soft, skin-safe silicone mixture (Wacker Elastosil Vario 15 with Catalyst Vario and silicone oil), allowing the device to contact objects and surfaces with skin-like compliance. This silicone layer surrounded a central rigid printed core that acted as the mechanical analogue of the finger’s bony structure. The inner surface that contacted the biological fingertip was padded with paint-roller foam to ensure comfort and to distribute pressure evenly across the distal phalanx. Standard elastics routed through printed tensioning elements provided passive return force, maintaining a stable coupling between finger and extension across different hand postures. Stainless steel screws were used for all fastening points, with press-fit nuts embedded directly in the printed material. The fingertip caps were additionally covered with soft Pritt Poster Buddies to prevent sharp edges and improve comfort during manipulation.

#### Dimensions and mass

Two device sizes were produced to accommodate hand size variability, with an additional medium thumb component used when needed. Ten participants used the small fingers with the small thumb, eight used the small fingers with the medium thumb, and two used the large fingers with the medium thumb. The standard 10 cm artificial fingers (without the thumb) weighed 116 grams, while the large version weighed 134 grams. The thumb components weighed 28 grams (small), 32 grams (medium), and 40 grams (large). The control (non-extending) devices were constructed with similar dimensions but were lighter: 60 grams for the standard fingers, 67 grams for the large fingers, and 11-15 grams for the thumb, depending on size. Thus, both devices added weight comparable to a small handheld tool, but crucially, the device was rigidly coupled to the finger rather than held in the hand.

#### Fitting and attachment

Each participant was individually fitted to ensure both stability and comfort. The fingertip caps were adjusted so that the distal phalanx was securely seated against the foam padding, and the elastic tension was tuned to maintain stable coupling when manipulating objects. Following fitting, participants performed a brief pick-and-place action with a small ball to confirm that the finger-extension device did not shift relative to the finger during movement. Participants were asked to report discomfort, and adjustments were made when necessary. No participant reported pain or loss of circulation during use.

#### Rationale for design in relation to proprioceptive mapping

The critical design property for the present study is the rigid and collinear coupling of the biological finger and the artificial finger segment. This contrasts with handheld tools, which typically permit slip, pivoting, or variable grasp posture. The finger-extension device therefore allowed us to treat the biological finger and the artificial finger as a single kinematic chain during proprioceptive mapping. Mapping spatial judgment along this chain permitted direct comparison of how biological and artificial segments are represented within the same coordinate frame, which was essential for testing whether and how the somatosensory system integrates artificial augmentation into body representation. A visual schematic of the finger-extension device design is provided in Supplementary Figure S1.

### Experimental design

Experiments 1A and 1B used a within-subject control design to isolate the effects of training with the finger-extension device on body representations. The same participants completed both sub-experiments (1A and 1B) on separate days (5–30 days apart), with the order of experiments counterbalanced across participants to mitigate between-day order effects. Each sub-experiment consisted of three parts: a touchscreen localization task, motor training, and a repetition of the localization task. A final motor task and questionnaire were also included but are not discussed in this article for brevity.

Within Experiment 1B, we deliberately trained with the short (1.4 cm) device but mapped with the long (10 cm) device both before and after training (see below). This design equates visual/proprioceptive context and total task exposure across stage 2 and stage 3 while manipulating only whether training involved true spatial augmentation. Participants were blindfolded during the brief swap to prevent visual recalibration of hand size.

### High-density proprioceptive localization task

The participants were seated comfortably in a cushioned chair with their left arm resting on a padded armrest. They were positioned at a table facing the short end of a 93.2 x 52.6 cm LCD touchscreen (iiyama ProLite TF4237MSC-B3AG, Tokyo, Japan), securely fixed above the table. The right arm was placed under the screen on a board with grooves for each finger to ensure consistent placement, and participants were instructed not to look under the screen. The screen displayed three vertical stripes corresponding to the right index, middle, and ring fingers.

### Localization task in the four task stages

#### Stage 1

The task of participants was to localize different locations on their right index finger using proprioception. The seven targets were evenly spaced between the base and fingertip along the finger’s midline. Throughout the task, the participants kept their right arm completely still and made localization judgments by touching the touchscreen with their left hand (Fig. S2). After the judgment was made, they were asked to move their left hand back to the resting start position, which was 16 cm outside the response field, adjacent to the screen edge.

On a given trial, the procedure was as follows. At the beginning of the trial, an image of a finger was displayed in the lower left corner of the screen. A red dot was placed at the target location within the finger. Participants then pointed on the screen to the corresponding target location on the stripe above their actual index finger. Participants then returned their left hand to the start position. No accuracy feedback was provided. This procedure was repeated for seven equally spaced locations across 20 trials, totaling 140 trials (see Fig. S3A for the layout of target locations). The order was randomized. This task took about 5 to 10 minutes depending on the participant’s speed.

#### Stage 2

The proprioceptive localization task was the same as in stage 1 with the exception that participants now also localized targets within the space of the biological finger and artificial finger (see Fig. S3B for the layout of the 14 target locations). There were seven distinct evenly spaced target locations within both the biological and artificial finger segments, for 14 distinct locations in total. There were again 20 trials per location, leading to 280 trials total. This task took around 10 to 15 minutes.

Stage 2 had the distinct procedure that it also required fitting the finger-extension device to the participant. After they finished the task in stage 1, they were asked to stay seated and remove their right arm from under the screen. The finger-extension device was fitted to their right hand for both experimental and control sessions. After making sure the finger-extension device fit comfortably, the participants’ right arm was placed under the screen again in the same position as before.

In Experiment 1A, the participants continued with the spatially augmenting finger-extension device. In Experiment 1B, after finishing both localization tasks, the finger-extension device was removed, and the control device was fitted in the same way as the finger-extension device.

#### Stage 3

After completing the motor tasks (see "Training protocol" below), participants were guided back to the touchscreen to repeat the localization task, which was the same proprioceptive localization task as in stage 2. In Experiment 1B, before starting this task, the control device was replaced with the finger-extension device while the participant was blindfolded. The blindfold ensured that participants did not see their hand without the finger-extension device, preventing visual input from recalibrating their hand representation and potentially diminishing the effects of the training. In Experiment 1A participants remained wearing the finger-extension device.

#### Stage 4

For the finger-only localization task, the device was removed while participants were blindfolded, and their hand was guided back under the screen before the blindfold was removed. Participants then performed the same proprioceptive localization task as in stage 1.

### Training protocol

The training took place between stages 2 and 3. During the training, participants were asked to complete a set of tasks to familiarize them with the use of the device (finger-extension or control) and train their motor control. They were instructed that the tasks should be completed predominantly with their right, device-wearing hand. That is, grasping and manipulation was to be performed solely with the right hand, and they were to only use their left hand in a supportive role when needed, that is, only to stabilize objects or assist minimally without performing the main action. This approach was chosen to make the training more ecologically relevant, as in daily life, both hands would typically be used to complete such tasks. Each task was repeated three times, forming three blocks of trials. Within each block, the order of tasks was consistent for each participant, but this order was randomized across participants.

The tasks included in the training were based on a previously introduced protocol (Kieliba et al., 2021): building a Jenga tower with wooden blocks, sorting plastic bricks according to their color, and sorting shapes by inserting them into a wooden box. In addition, participants completed the Purdue Pegboard Test (both long and short peg versions) before and after the three repetitions (Lawson, 2019). See the Supplementary Information for a detailed description of each task.

### Data analysis

The data were tested for normality using the Shapiro–Wilk test and found to be normally distributed (p > .05); therefore, parametric statistics were used. Linear regression was used to quantify the represented length of finger and artificial finger segment. In the localization tasks, the perceived location of each target was calculated as the mean of participants’ response distributions. The true position of the targets was determined from participants’ finger measurements and the known device length. By performing regressions of perceived versus true locations, we obtained an offset (β_0_) and a slope (β_1_). The offset was discarded, as it only reflected a spatial bias without providing information on length perception. The slope, however, indicates the scaling factor and represents participants’ length perception.

For the finger-only localization task, one regression was performed per condition (pre- and post-training), resulting in four slopes per participant (two per sub-experiment). For the finger+extension localization task, two regressions were performed per condition (pre- and post-training): one for the biological finger segment and one for the artificial finger segment, yielding eight slopes per participant. These slopes quantified perceived spatial scaling along each segment and served as a measure of perceived length.

To directly test whether training with the finger-extension device selectively increased overall represented length relative to training with the non-augmenting control, we also computed a combined bio+artificial slope at Stages 2 and 3. For this measure, we pooled the 14 targets (7 biological + 7 artificial) and fit a single linear regression of perceived versus true location across the full bio+artificial surface for each participant and sub-experiment. These combined slopes were submitted to a 2 x 2 repeated-measures ANOVA with within-subject factors Stage (2 vs. 3) and Device (finger-extension [Experiment 1A] vs. control [Experiment 1B]).

The comparison of the represented length otherwise involved analyzing the segments separately, leading to two comparisons of condition and pre/post-training. Changes in slope across experimental stages (e.g., before and after training) were evaluated using paired t-tests. We conducted planned comparisons between stage 1 vs. 2 (biological finger), 2 vs. 3 (biological finger and artificial finger), 1 vs. 4 (biological finger), 3 vs. 4 (finger) within each of Experiment 1A and 1B, resulting in five paired t-tests per experiment. Reported *p*-values were Bonferroni-corrected within each experiment to account for five planned comparisons.

All *p*-values are two-tailed and are reported alongside 95% confidence intervals and Bayes Factors (BFs). For Bayesian analyses, the default Cauchy prior was used.

## Acknowledgments

We would like to thank Sibrecht Bouwstra for her key role in the development of the finger-extension devices, as well as the Technical Support Group of the Donders Centre for Cognition for their additional support.

## Author contributions

Conceptualization: DR, SG, VCP, LEM; Methodology: DR, SG, VCP, LEM; Investigation: SG; Visualization: DR, SG; Supervision: VCP, LEM; Writing—original draft: DR, LEM; Writing—review & editing: DR, SG, VCP, LEM.

## Data and materials availability

The data and code that support the findings of this study will be available from the Open Science Framework upon publication (https://osf.io/wpshu/).

