## Supplementary File for "Dynamics of sensorimotor plasticity during spatial finger augmentation"

Dominika Radziun* *et al.*

**This PDF file includes:**

Structural components of the finger-extension device
Experimental setup for localization task
Training task descriptions and outcome measures
Supplementary videos

**Structural components of the finger-extension device**

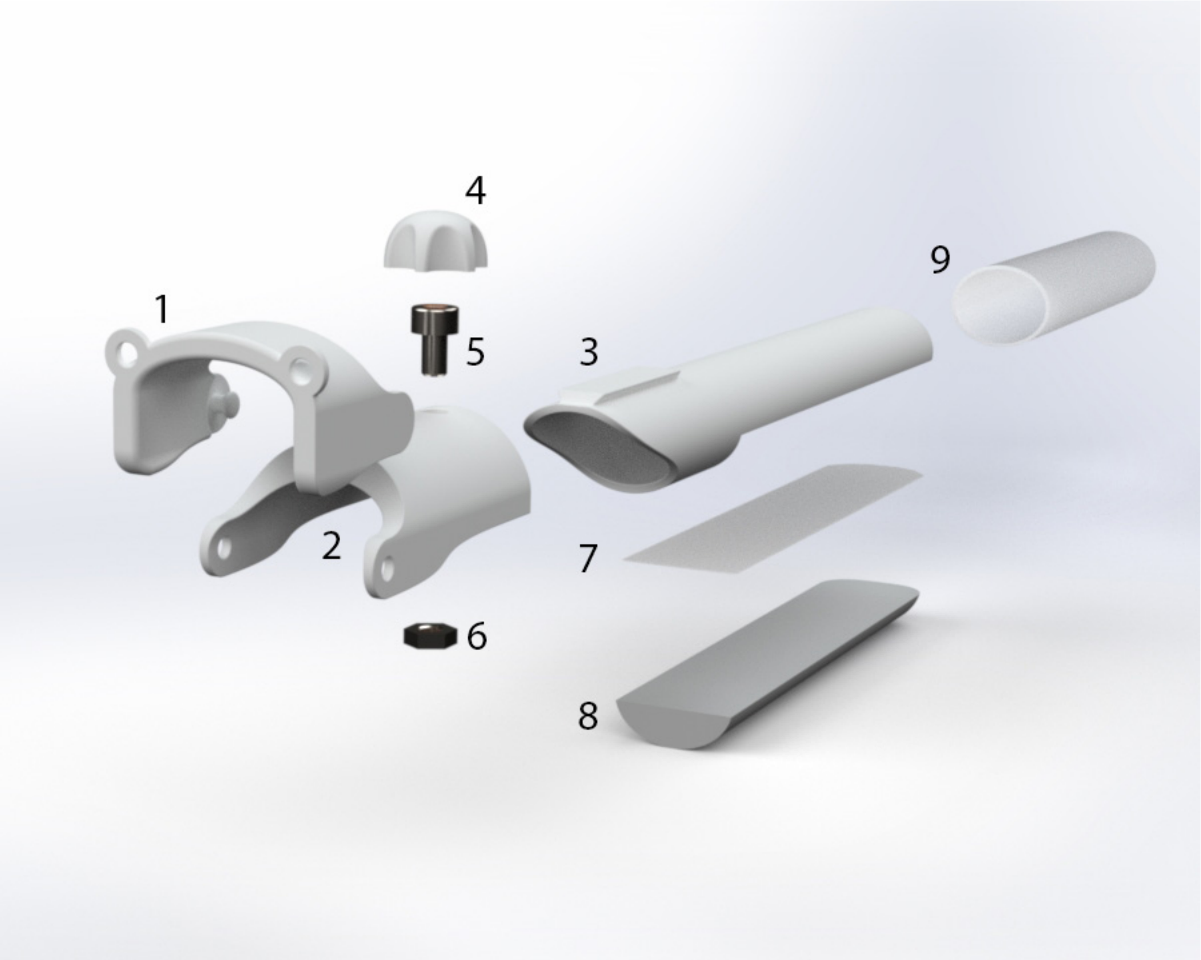


**Figure S1.** (1) Proximal segment with joint and elastic-band connection. (2) Intermediate segment with joint and adjustable length to the fingertip. (3) Fingertip cap. (4) Knob for securing the fingertip cap. (5) Fastening screw for fingertip attachment. (6) Embedded nut for fingertip fastening. (7) Double-sided tape used to secure the internal padding. (8) Foam padding providing a comfortable contact surface with the biological fingertip. (9) Silicone sleeve forming the external gripping surface.

### Experimental setup for localization task

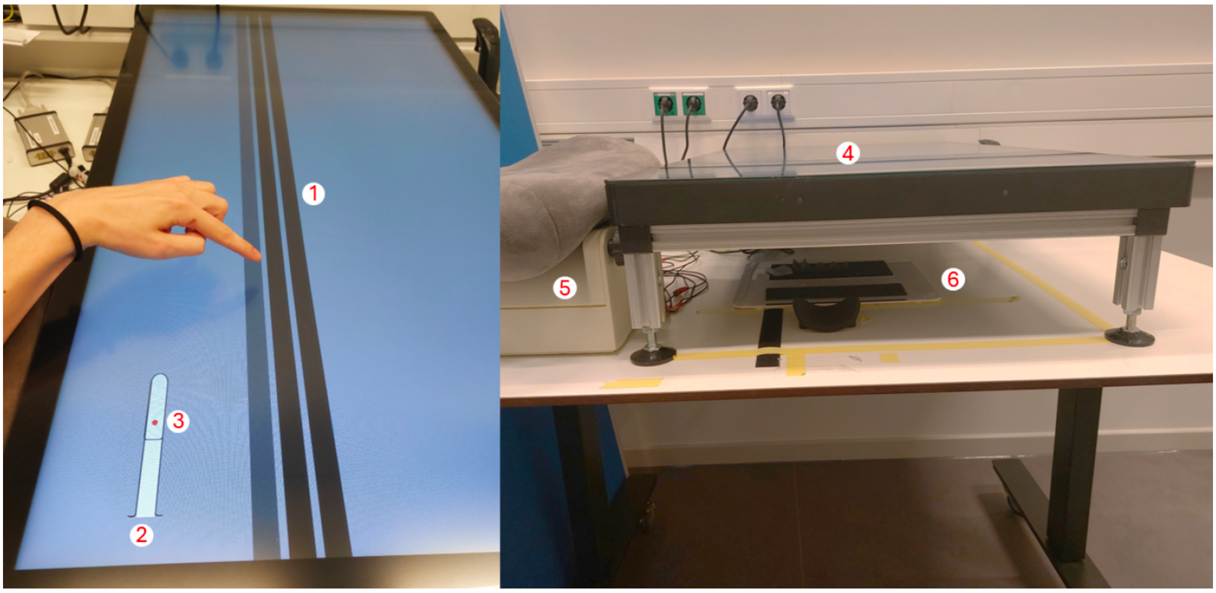


**Figure S2.** A top-down view of the finger+extension localization task (left) and a frontal view of the localization task setup (right). (1) The three stripes are aligned with the index, middle, and ring fingers placed under the touchscreen. The left stripe is colored in gray to indicate the finger of interest. (2) The image of a finger with the finger-extension device shown to the participant on which the requested locations are displayed. (3) One of the 14 requested locations displayed on the artificial finger in red. (4) The elevated touchscreen under which the right hand is placed. (5) The elevated armrest on which the right elbow is placed. (6) The board with the finger grooves on which the right hand is placed together with a small armrest for the forearm.

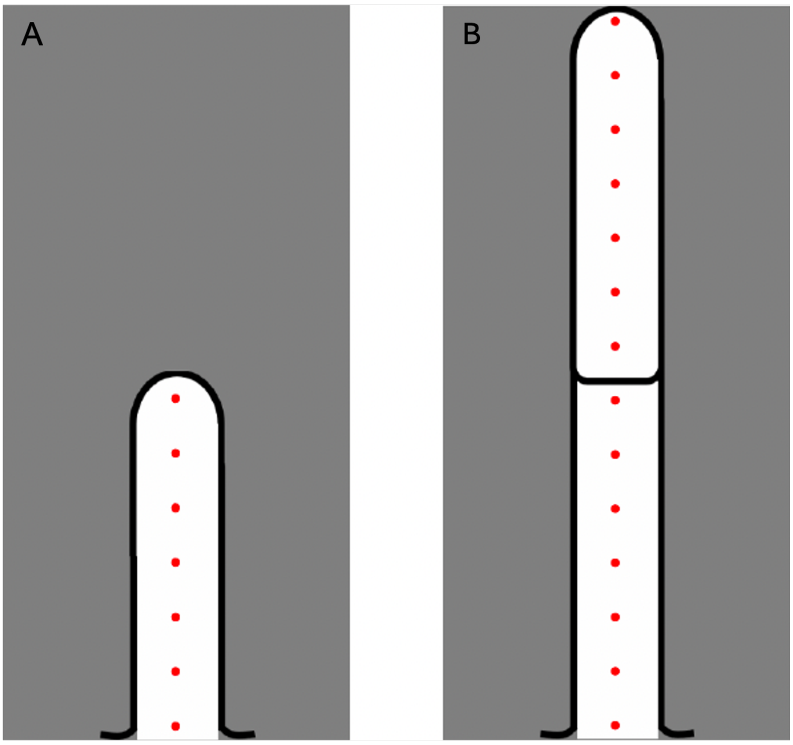


**Figure S3.** The distribution of 7 locations for the finger (A) and 14 locations for the finger+extension (B) localization tasks.

### Training task descriptions and outcome measures

Following the 30–45 minutes training period, participants' performance with the finger-extension device was assessed across multiple tasks. Detailed statistical analyses confirmed a significant learning effect over time.

Task completion times (the primary outcome measure) were assessed for normality using the Shapiro–Wilk test. When any repetition within a task significantly deviated from normality, the entire task was analyzed using a non-parametric method (Wilcoxon signed-rank test); otherwise, parametric methods were applied. Planned pairwise comparisons were conducted between blocks (block 1 vs. block 2, block 2 vs. block 3) using two-tailed tests. Statistical comparisons involving were evaluated against a Bonferroni-adjusted significance threshold of $\alpha = 0.025$ to account for multiple planned comparisons.

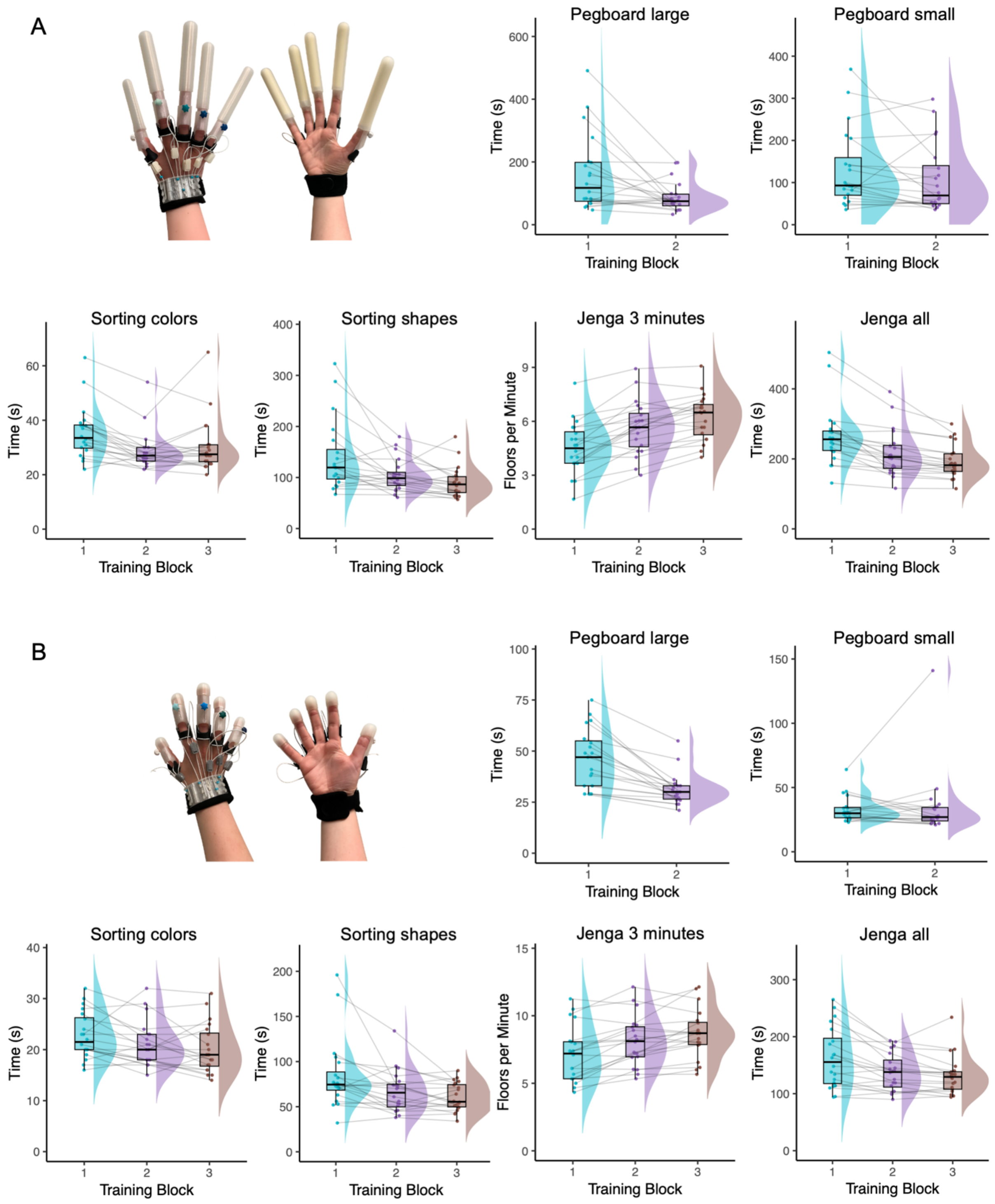


**Figure S4.** Overview of training tasks performed in Experiment 1A (A) and Experiment 1B (B). In Experiment 1A, participants trained with a finger-extension device; in Experiment 1B, participants trained with a control device. Each panel displays two rows of task conditions: the top row shows the pegboard transfer tasks with large and small pegs, and the bottom row shows the sorting colors task, sorting shapes task, the Jenga 3-minute task, and the Jenga all task.

**Pegboard transfer task**

Before and after the training, participants completed a custom task inspired by the Purdue pegboard task. The task was divided into two stages: one involving larger pegs (9 cm long) and one involving smaller pegs (5 cm long), both with a diameter of 0.7 cm at the ends and 0.8 cm at the midpoint. All pegs were made from standard PVC stock and lathed at the ends for improved grip. Participants were asked to transfer 14 pegs from one wooden pegboard to another, using only their right hand, which was fitted with the finger-extension device. The pegboard featured 27 holes arranged in 9 rows and 3 columns. Grasping and manipulation were to be performed solely with the right hand, while the left hand was allowed only to stabilize the pegboard. The outcome measure was the time taken to complete the transfer.

In Experiment 1A, which involved training with the finger-extension device, participants showed significant improvements in the large peg condition. Completion times were significantly reduced following the training session (Wilcoxon $V(19) = 27.0$, $p = .006$, $BF_{10} = 5.67$), indicating improved gross motor performance. Performance in the small peg condition did not improve significantly (Wilcoxon $V(19) = 59.0$, $p = .09$, $BF_{10} = 0.56$).

In Experiment 1B, which involved training with a control device, participants significantly improved their performance in the large peg condition (Wilcoxon $V(18) = 0.0$, $p < .001$, $BF_{10} = 2185.93$). No significant change was found in the small peg condition (Wilcoxon $V(18) = 56.5$, $p = .206$, $BF_{10} = 0.25$). Excluding one outlier due to a technical issue with the coating on the device's fingertips did not alter the outcome (Wilcoxon $V(17) = 38.5$, $p = .13$, $BF_{10} = 0.42$). Due to a technical issue that completely prevented one participant from completing the task in the post-training session, the initial total number of participants for this analysis was 19 instead of 20.

**Sorting colors task**

In the *sorting colors task*, participants were asked to sort LEGO® DUPLO® bricks into plastic trays according to color, using only their right hand, which was equipped with the finger-extension device. The task involved four colors, yellow, red, green, and blue, and included a fixed number of bricks in two standard sizes. The set comprised two large yellow bricks, one small yellow brick, three large red bricks, six small red bricks, one large green brick, two small green bricks, one large blue brick, and three small blue bricks. Small bricks measured 32 mm × 32 mm in width and depth, with a height of 19.2 mm (22 mm including studs), while the large bricks were 32 mm wide, 64 mm long, and approximately the same height. The outcome measure was the time required to complete the task.

In Experiment 1A, participants trained with the finger-extension device showed a significant improvement in sorting performance following the initial training session. Task completion time decreased significantly from session 1 to session 2 (Wilcoxon $V(19) = 8.5$, $p = .006$, $BF_{10} = 2782.66$), indicating a rapid learning effect. However, no further improvement was observed between session 2 and session 3 (Wilcoxon $V(19) = 61.5$, $p = .95$, $BF_{10} = 0.397$), suggesting that most of the learning occurred early in the training and plateaued thereafter.

In Experiment 1B, participants using the control device did not show statistically significant improvement between session 1 and session 2 (Wilcoxon $V(19) = 31.0$, $p = .06$, $BF_{10} = 1.24$), nor between session 2 and session 3 (Wilcoxon $V(19) = 49.0$, $p = .38$, $BF_{10} = 0.47$). This indicates that training with the control device did not lead to measurable performance enhancement across sessions, in contrast to the group using the finger-extension device.

**Sorting shapes task**

In the *sorting shapes task*, participants were asked to insert wooden blocks into their corresponding holes in a wooden shape-sorting box. The set included 16 uniquely shaped and colored blocks: a yellow rhombus, yellow trapezoid, yellow pentagon, red triangle, red trapezoid, red hexagon, green square, green rectangle, green octagon, blue oval, blue star, and blue flower. Each shape was represented once. Participants were instructed to carry out all grasping and manipulation with their right, device-wearing hand, while the left hand could be used only to support or reposition the box if needed. The outcome measure was the time taken to insert all shapes correctly into their respective slots.

In Experiment 1A, involving the finger-extension device, participants demonstrated significant improvements in task performance across sessions. Completion time decreased significantly from session 1 to session 2 (Wilcoxon $V(19) = 32.0$, $p = .01$, $BF_{10} = 5.52$), suggesting that the device supported the development of improved control and efficiency during shape manipulation. A further improvement was observed from session 2 to session 3 (Wilcoxon $V(19) = 36.0$, $p = .036$, $BF_{10} = 1.50$), indicating continued performance gains with additional practice.

In Experiment 1B, where participants trained with the control device, significant improvement was observed between session 1 and session 2 (Wilcoxon $t(19) = 29.5$, $p = .01$, $BF_{10} = 3.70$). However, no further improvement occurred from session 2 to session 3 (Wilcoxon $t(19) = 66.0$, $p = .31$, $BF_{10} = 0.74$). This pattern indicates that while early gains occurred in both experimental contexts, ongoing improvements were specific to training with the finger-extension device.

**Jenga 3-minute task**

In the *Jenga 3-minute task*, participants were instructed to build as many complete floors of a Jenga tower as possible within a fixed time limit of three minutes. Each floor consisted of three standard rectangular wooden blocks placed side by side. Participants used only their right hand, equipped with the finger-extension device, to manipulate and place the blocks. The left hand was not permitted during the task. The primary outcome measure was the number of full floors completed within the three-minute window.

In Experiment 1A, where participants trained with the finger-extension device, performance improved significantly across sessions. The number of completed floors increased from session 1 to session 2 ($t(19) = -5.98$, $p < .001$, $BF_{10} = 2343.80$), and further gains were observed from session 2 to session 3 ($t(19) = -4.31$, $p < .001$, $BF_{10} = 86.28$). These results suggest that participants became progressively more efficient at using the device for the task, benefiting from both initial learning and continued refinement over multiple training blocks.

In Experiment 1B, conducted with the control device, participants also showed a significant improvement between session 1 and session 2 ($t(19) = -4.70$, $p < .001$, $BF_{10} = 188.06$). However, no statistically significant improvement was observed between session 2 and session 3 ($t(19) = -2.43$, $p = .05$, $BF_{10} = 2.40$). This pattern shows that sustained gains across sessions were more pronounced in the context of training with the finger-extension device.

**Jenga all task**

In the *Jenga all task*, participants were asked to build a complete Jenga tower consisting of 18 floors, with each floor made up of three wooden blocks. Unlike the 3-minute version, this task required the tower to be completed in its entirety, regardless of how long it took. As in the other tasks, participants were instructed to use only their right hand, which was fitted with the finger-extension device, to manipulate and place the blocks. The left hand was not permitted. The outcome measure was the total time required to complete the full tower.

In Experiment 1A, which involved training with the finger-extension device, participants showed significant improvement across sessions. Completion times decreased between session 1 and session 2 (Wilcoxon $t(19) = 1.0$, $p < .001$, $BF_{10} = 974.07$), and continued to improve from session 2 to session 3 (Wilcoxon $t(19) = 8.0$, $p < .001$, $BF_{10} = 30.45$). These results indicate consistent gains in efficiency when using the finger extensions, suggesting ongoing refinement with repeated use of the device.

In Experiment 1B, when participants performed the training using the control device, a significant reduction in completion time was also observed from session 1 to session 2 (Wilcoxon $t(19) = 22.5$, $p = .002$, $BF_{10} = 45.24$). However, no significant improvement was found between session 2 and session 3 (Wilcoxon $t(19) = 51.0$, $p = .08$, $BF_{10} = 0.68$). As with the 3-minute task, this suggests that while initial exposure to the task led to performance gains under both experimental contexts, continued improvement was more robust and sustained in the presence of active device support.

### Supplementary videos

The individual appearing in the supplementary videos provided explicit informed consent for their recordings to be captured and publicly shared as part of this article.

**Supplementary Video 1.** Demonstration of the motor training tasks performed with the finger-extension devices (sorting pegs into holes, sorting colored blocks, stacking a Jenga tower). Available at: https://osf.io/wpshu/files/4dawg.

**Supplementary Video 2.** Demonstration of fitting the finger-extension device.
Available at: https://osf.io/wpshu/files/6sqw9.
